# devider: long-read reconstruction of many diverse haplotypes

**DOI:** 10.1101/2024.11.05.621838

**Authors:** Jim Shaw, Christina Boucher, Yun William Yu, Noelle Noyes, Heng Li

**Affiliations:** Department of Data Science, Dana-Farber Cancer Institute, Boston, MA 02215, USA; Department of Biomedical Informatics, Harvard Medical School, Boston, MA 02215, USA; Department of Computer and Information Science and Engineering, University of Florida, Gainesville, FL 32611, USA; Ray and Stephanie Lane Computational Biology Department, Carnegie Mellon University, Pittsburgh, PA 15213, USA; Department of Veterinary Population Medicine, University of Minnesota, St. Paul, MN 55421, USA

**Keywords:** long-reads, haplotyping, viruses, genes, metagenome, de Bruijn graph

## Abstract

Reconstructing haplotypes is important when sequencing a mixture of similar sequences. Long-read sequencing can connect distant alleles to disentangle similar haplotypes, but handling sequencing errors requires specialized techniques. We present `devider`, an algorithm for haplotyping small sequences—such as viruses or genes—from long-read sequencing. `devider` uses a positional de Bruijn graph with sequence-to-graph alignment on an alphabet of informative alleles to provide a fast assembly-inspired approach compatible with various long-read sequencing technologies. On a synthetic Nanopore dataset containing seven HIV strains, `devider` recovered 97% of the haplotype content compared to 86% for the next best method while taking < 4 minutes and 1 GB of memory for > 8000× coverage. Benchmarking on synthetic mixtures of antimicrobial resistance (AMR) genes showed that `devider` recovered 83% of haplotypes, 23 percentage points higher than the next best method. On real PacBio and Nanopore datasets, `devider` recapitulates previously known results in seconds, disentangling a bacterial community with > 10 strains and an HIV-1 co-infection dataset. We used `devider` to investigate the within-host diversity of a long-read bovine gut metagenome enriched for AMR genes, discovering 13 distinct haplotypes for a tet(Q) tetracycline resistance gene with > 18, 000× coverage and 6 haplotypes for a CfxA2 beta-lactamase gene. We found clear recombination blocks for these AMR gene haplotypes, showcasing `devider`’s ability to unveil ecological signals for heterogeneous mixtures.

## 1 Introduction

The presence of highly similar genomic sequences within a single or a group of organisms is common in biological settings. Examples include viral quasispecies [1] in single-stranded RNA virus populations (e.g., HIV-1 and SARS-CoV-2) [2] or co-existing microbial subspecies in microbiomes [3]. Small genomic differences can have large functional implications [4,5], so it is crucial to disentangle this heterogeneity. We will call the process of recovering similar genomic sequences “haplotyping” or “phasing”. While traditionally used in the context of diploid organisms, we extend the concept here to encompass the resolution of genetic diversity in microbes, viruses, or even genes.

With high-throughput sequencing, we can obtain haplotypes by linking reads that share informative alleles, for example, single nucleotide polymorphisms (SNPs). Unfortunately, standard de novo short-read or nanopore long-read assembly approaches can collapse small-scale variation [6], returning only a consensus sequence. Although haplotype-resolved assembly has become standard for PacBio HiFi sequencing [7,8,9,10], HiFi data may not always be available and assembly is computationally intensive. In contrast to assembly approaches, reference-based haplotyping uses a reference plus alignment to facilitate haplotyping; many existing approaches use the alignment, SNP calling, then phasing paradigm [11,12,13,14,15,16,17,18,19].

We are interested in reference-based haplotyping for (1) long-read sequencing, (2) small sequences of approximately the read length, and (3) an *unknown*, possibly large number of haplotypes. Long reads can connect more distant alleles across shared genomic regions than short reads. Still, a technical challenge is dealing with sequencing errors for certain technologies, e.g., Oxford Nanopore reads can have 90™99% sequencing accuracy depending on the chemistry and basecalling [20]. We focus on small sequences on the order of the read length (i.e., “local haplotyping” [21]), but we do not necessarily require all reads to overlap the region of interest. This is sufficient for haplotyping genes of interest or estimating diversity. While this seems like a simple task, systematic errors, high coverage, and low abundance haplotypes make accurate reconstruction challenging [22].

Many haplotyping algorithms work for either long reads or an unknown number of haplotypes, but only a few were designed to do both. RVHaplo [13], CliqueSNV [12], and iGDA [11] are long-read methods that call low-frequency SNPs and can phase diverse genomic sequences; RVHaplo uses a network clustering formulation, CliqueSNV uses a clique-merging approach, and iGDA uses a probabilistic local haplotyping step with an overlap-layout algorithm for reconstruction. Another class of approaches focuses on genome-scale haplotype reconstruction for prokaryotes [23,24,25], however, these methods have not been tested for phasing high-diversity communities with > 5 similar haplotypes.

We present `devider`, a new long-read, reference-based haplotyping method for diverse small sequences. Given a set of aligned reads and SNPs, `devider` models the haplotyping problem as an assembly problem on a positional de Bruijn graph (PDBG). `devider` is inspired by the kSNP algorithm [18] which similarly uses a PDBG but for haplotyping only diploids. We use the fact that the PDBG naturally splits if enough variation is present and collapses under ambiguity, thus haplotyping samples without prior knowledge of the number of distinct sequences. We then leverage the length of long reads by finding walks along the PDBG through unitig construction and read-to-graph alignment. We find that `devider` efficiently resolves haplotypes on a variety of synthetic and real datasets and show its versatility for unveiling genomic heterogeneity.

## 2 Methods

At a high level, `devider` takes a set of aligned reads in BAM format plus SNPs in VCF format. It then outputs the sequences of SNPs for each recovered sequence (i.e., the haplotype), the abundance of each haplotype, and a base-level sequence for each haplotype. `devider` does not call SNPs or perform alignment, but we supply a wrapper for `devider` with minimap2 [26] and LoFreq [27] for alignment and SNP calling. We use LoFreq for variant calling unless otherwise stated, but users can choose to use their own SNP caller.

`devider` follows in three major steps (**Fig. 1 a-c**): (1) encoding the aligned reads with SNPs, (2) constructing a PDBG and positional unitig graph while filtering errors, and (3) aligning the SNP-encoded reads to the positional unitig graph to obtain walks along the graph that are supported by many reads, which represent candidate haplotypes.

**Fig. 1.**
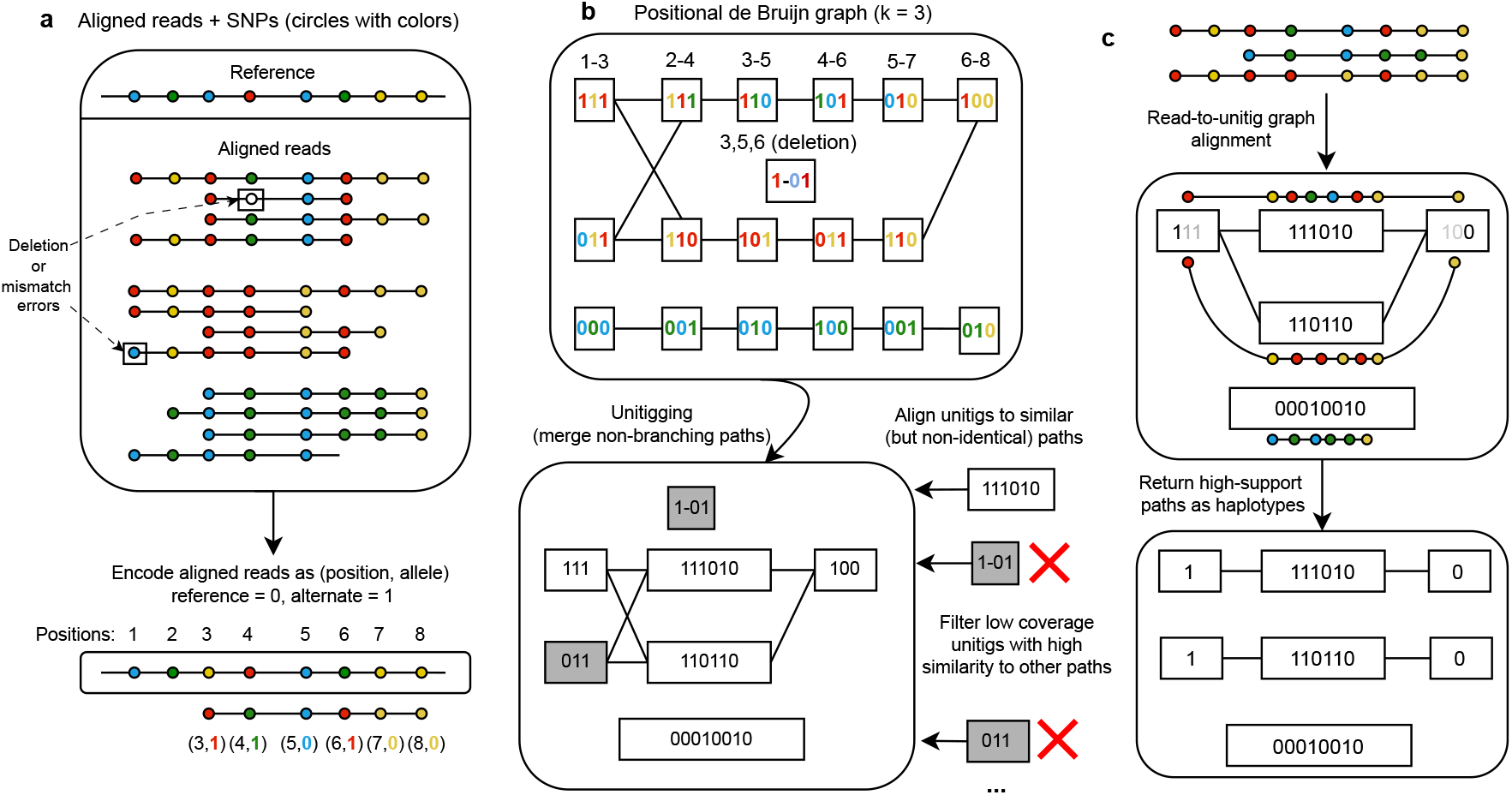
Algorithmic framework for `devider`. **a**. Reads that are aligned to a reference are converted to a SNP representation with positional information. Sequencing errors lead to erroneous SNP encodings. **b**. The SNP-encoded reads are turned into a positional de Bruijn graph (PDBG) (k = 3 shown). In a PDBG, k-mers are collapsed if their alleles *and* their positions are identical. Errors in reads lead to spurious k-mers in the PDBG. After merging paths with in-degree and out-degree equal to 1 (unitigging), unitigs are aligned back to the graph to filter low-coverage, high-similarity unitigs. **c**. Reads are aligned back to the filtered unitig graph to determine high-confidence walks through the graph. These paths are taken to be putative haplotypes. `devider` then post-processes the haplotypes to output haplotype abundances, a base-level consensus of each haplotype, and the reads assigned to each haplotype.

### 2.1 Encoding reads with informative SNPs and filtering false positives

The first step is to filter false positive SNPs arising from strand-specific errors. For each SNP, we apply a simple strand bias filter by using Fisher’s exact test [28] on the 2-by-2 contingency table defined by forward/reverse strands and reference/alternate alleles. For high coverage, Fisher’s exact test is too stringent, so we also require an odds ratio of > 1.5 or < 1*/*1.5 for the contingency table. We then apply the Benjamini-Hochberg [29] multiple-testing correction at 0.005 FDR threshold. More powerful methods are possible for strand filtering [30], but we opted for a simple method because we are only concerned with phasing.

We then subsample the SNPs if too many are present (i.e., the sequences are highly diverged) as follows: for a given parameter *α* (discussed in Section 2.5) and median number of SNPs contained in a read *β*, we downsample uniformly to retain only *α/β* of the SNPs if *α < β*. We do this because when too many SNPs are present, a group of SNPs may span only a small region of the genome, which we wish to avoid in our subsequent SNP-based *k*-mer approach. We then realign each read using blockaligner [31] against the 32 bp flanks around each filtered SNP site, replacing the site with all possible alleles and then selecting the allele that gives the highest alignment score.

Finally, we encode each read as an ordered list of tuples such as (3, 1), (4, 1), (5, 0), (6, 1), (7, 0), (8,0); see **Fig. 1 a**. The first number represents the *i*-th SNP in the reference the read is aligned to, and the second number indicates the allele that the read contains where 0 is reference and 1 to 3 indicate alternates. We skip SNPs if either the SNP site has a deletion in the read or the read’s base at the SNP site is neither an alternate nor reference allele in the VCF. Thus, a sequence such as (3, 1), (5, 0) is possible.

### 2.2 Positional de Bruijn graph and unitigs on the SNP alphabet

All reads will be considered SNP-encoded reads from now on. A SNP-encoded read is a string on an alphabet *Σ* = 𝕫^+^×{ 0, 1, 2, 3} subject to the constraint that the first symbol in each letter is in increasing order, e.g, (5,1) must come before (7,0). *k*-mers of SNP-encoded reads are defined as elements in Σ^*k*^, and we construct a de Bruijn graph in the usual manner by collecting all *k*-mers within the reads and adding directed edges between *k*-mers that overlap *k* ‒ 1 SNPs (**Fig. 1b**, top). However, because the positions are encoded in the alphabet Σ, we only collapse k-mers if they have the same positions. This is now a *positional* de Bruijn graph (PDBG) [32,33,34]. The PDBG is a directed acyclic graph (DAG) since any cycle would violate the increasing ordering for the SNPs. This fact will be important for the subsequent sequence-to-graph alignment steps.

We automatically choose a value of *k* as follows. Let *γ* be the 33rd percentile of the number of SNPs contained in a read and let *N* be the number of SNPs in the reference (after filtering). Let *M* be a parameter representing the maximum possible value of *k*, which is set to avoid long error-prone *k*-mers. This is dependent on the sequencing technology and is picked through a preset option. We let 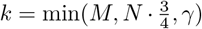. We do not let *k* span more than 75% of the reference to not miss *k*-mers if a smaller haplotype only covers a subsection of the reference. We discuss parameter choices of *M* in Section 2.5 and show results over varying values of *k* in Section 3.1.

We apply an initial filtering step to discard likely erroneous *k*-mers. Let the coverage of a *k*-mer be the number of times it appears in a SNP-encoded read, and *m* be the mean *k*-mer coverage. Let *A* be the minimum allowable abundance for a haplotype (default = 0.0025 or 0.25%). We filter *k*-mers that appear only once or have less than *m* ·*A* coverage. Finally, from the filtered PDBG, we construct the positional unitig graph by merging all non-branching paths into unitigs (**Fig. 1b**, bottom) [35]. We let the coverage of the unitig be the mean coverage of the merged k-mers.

### 2.3 Filtering unitigs by error-aware unitig-to-graph alignment

The main technical challenge is to simplify the unitig graph, which can still have many spurious unitigs arising from *k*-mer errors (**Supplementary Figure 1 and 2**). In standard de Bruijn graph assembly, tip removal and bubble popping are used to remove noise *and* variation [36]. However, we do not want to remove true variation. Thus, we use a unitig-to-graph alignment approach plus coverage information for fine-grained unitig filtering; this generalizes both tip removal and bubble popping but uses alignment information and coverage information.

#### Classifying errors

Let *G* be the positional unitig graph. Recall that a unitig node *v* can be represented by *v* = (*x*_1_, *x*_2_, …, *x*_*n*_) where *x*_*i*_ = (*a*_*i*_, *b*_*i*_) with *a*_*i*_ the SNP position and *b*_*i*_ the allele. Given two unitigs *v*_1_ and *v*_2_, we classify errors as SNP deletions (del), reference-to-alternate (rtoa) mismatches, and alternate-to-reference (ator) mismatches as follows.

- *s*(*v*_1_, *v*_2_) is the number of SNPs that are the same in *v*_1_ and *v*_2_ (i.e., share the same position and base).
- *del*(*v*_1_, *v*_2_) is the number of SNPs in *v*_1_ are deleted relative to *v*_2_ (i.e., do not appear in *v*_2_ and lie between the first and last SNP of *v*_2_).
- *rtoa*(*v*_1_, *v*_2_) and *ator*(*v*_1_, *v*_2_) represent the number of SNPs that have the same position but different alleles between *v*_1_ and *v*_2_. Specifically, *rtoa*(*v*_1_, *v*_2_) is the number of differing SNPs for which *v*_1_ has the *reference* allele (and thus *v*_2_ has an alternate allele), whereas *ator* is the number of differing SNPs for which *v*_1_ has an *alternate* allele.

We stratify the error types because they appear with different frequencies. *rtoa* and *ator* errors differ because of reference bias: if a read comes from a haplotype with a true alternate allele, it may systematically align to the reference allele incorrectly [37]. A SNP deletion (*del*) error is the most common due to the following reason: consider a biallelic site with two alleles, (A, C) and a read originating from the haplotype with allele A. `devider`’s convention is to consider the read’s SNP to be deleted if there is a base-level deletion in the CIGAR string or the read’s base is G or T at the SNP site. Of the four error possibilities (deletion, substitution to C, substitution to G, substitution to T), three of them result in a SNP deletion in the SNP-encoded reads. Furthermore, long reads can have inherently higher deletion error rates as well [38].

#### Alignment to DAG

Next, we align the unitigs back to the unitig graph to find an alignment path. However, we disallow alignments back to unitig itself since this would always be the best match. If there is a path such that *s* (successes) is large relative to *del, rtoa* and *ator* (errors) and this path has much higher coverage than the unitig, then the unitig is likely an error that originates from the path.

By abuse-of-notation, let *s*(*v, P*) refer to the number of matching alleles between *v* and the string spelled out by a unitig path *P*. We wish to find a path that does not contain *v* and maximizes *s*(*v, P*). Furthermore, we impose that (1) ties are broken by taking the highest coverage path, (2) *v*’s first and last SNP positions must be within *P*‘s first and last SNP positions, and (3) all unitigs in *P* overlap *v*. We do not need to penalize for the error terms in this step because more matching alleles imply fewer deletions and errors.

Due to the DAG structure of the PDBG (and thus the positional *unitig* graph), the optimal path can be found with standard dynamic programming. Consider a unitig *v* and a topological order on the nodes 1, …, *n* that overlap *v*, we wish to find the optimal path ending at node *v*_*j*_. Let *P*_*j*_ be a path that ends at node *v*_*j*_. The following recurrence holds:

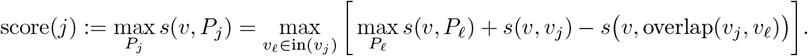

Note that we subtract the overlap to avoid double counting, since *v*_*j*_ and its incoming unitigs may overlap. After obtaining all scores, we find optimal paths that satisfy the three constraints above. Each alignment takes *O*(|*V*| |*E*|) worst-case running time, but in practice, unitig graphs are sparse (e.g. **Supplementary Fig. 1**), so this step is not a bottleneck.

#### Error-aware filtering

Once we have a best path *P* for each unitig *v*, we use an error-aware, one-sided Binomial test based on coverage to filter spurious unitigs. We run the alignment and error-aware filter starting from the smallest coverage unitig, then remove unitigs from the graph, and repeat for the next smallest coverage unitig.

Let *cov*(*v*) be the unitig coverage, and let *cov*(*P*) be either (1) the highest coverage unitig in *P* that completely covers *v* or (2) the mean unitig coverage in *P* if no unitigs in *P* completely cover *v*. We filter out *v* if Pr(*cov*(*v*) ≥*Binomial*(*cov*(*P*), *q*)) < 0.005, where *q* is defined as follows: define *p*_*del*_ = 0.35, *p*_*rtoa*_ = 0.15, and *p*_*ator*_ = 0.10 to model the error frequency behavior as previously discussed. Then 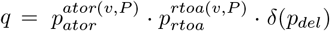 where δ(*p*_*del*_) = *p*_*del*_ if *ator*(*v, P*) + *rtoa*(*v, P*) = 0 but otherwise equal to 1. This formula slightly deviates from the independence assumption of the three error modalities. We found that systematic biases could occasionally cause two errors to occur non-independently. Thus, we loosen the independence on *p*(*del*), which was the main cause of errors. Overall, we found that these values are conservative and work for error-prone reads (≈ 95%), but could likely be tightened to improve sensitivity in the future as sequencing technologies improve.

### 2.4 Read-to-graph alignment and collecting haplotype paths

To find candidate haplotype paths through the filtered unitigs, we align reads back to the graph using a sequence-to-DAG alignment algorithm (**Fig. 1 c**), which is similar to unitig-to-graph alignment. We will use these alignments to find well-supported walks through the PDBG, representing candidate haplotypes.

Let *r* be a read. Here, rather than finding a path to maximize *s*(*r, P*) as we did before, we find a path maximizing *s*(*r, P*) ‒ 3× *err*(*r, P*) where *err* = *ator*(*r, P*) + *rtoa*(*r, P*). The penalty term of 3 was chosen to penalize errors more than similarities. Although true deleted SNPs in a haplotype are possible, we assume deletions mainly occur due to noise, so we do not penalize deletions. Compared to unitig-to-graph alignment, we add two changes. First, we allow *P* to bridge sinks-to-sources. This allows the alignment to rescue broken paths due to erroneous filtering. Secondly, we do not require the path to completely cover the read, also in case of erroneous filtering. We use the same dynamic programming procedure as for unitig-to-graph alignment; *err*(*r, P*) can also be split into the exact same recurrence and is thus solvable by dynamic programming.

After aligning all reads, we have a set of paths with read-alignment multiplicities. If two or more equally good paths exist for a read, we do not assign the read to any path. If a path is contained within another, we remove the contained path. If the path is *uniquely* contained in another, we add the contained path’s multiplicity to the non-contained path’s multiplicity. We filter for high-confidence paths by removing paths for which the read-alignment multiplicity is < 3× the minimum unitig coverage within the path.

#### Haplotype consensus and outputs

For the filtered paths, we assign the reads to each path by finding the path that maximizes 2 \**s*(*r, P*) ‒ 3\**rtoa*(*r, P*) ‒ 5 \**ator*(*r, P*) ‒ *del*(*r, P*), assigning the read to no path if the score is less than 0. The penalty weights are heuristically chosen to reflect the frequency of occurrence, e.g., deletions are common so they are penalized less. We then take the consensus allele for each SNP site after assignment and the abundance as the fraction of assigned reads.

Lastly, we perform a deduplication step as follows. For some resolution parameter *ρ* (see Section 2.5), we merge two resulting consensus haplotypes *H*_1_, *H*_2_ together if *rtoa*^Δ^(*H*_1_, *H*_2_)+*ator*^Δ^(*H*_1_, *H*_2_) */s*^Δ^(*H*_1_, *H*_2_) *< ρ*, where the Δ considers only unambiguous SNPs for which > 0.75 fraction of reads carry the majority allele. After merging, we reassign the reads to the best candidate haplotype subject to the same scoring scheme above, take consensuses, and filter low-abundance and low-depth haplotypes (default = 0.25% and 5× depth). We iterate this procedure until the number of haplotypes does not change. We then return the abundances, the reads assigned to each haplotype, and the sequences of the SNPs for each haplotype. Finally, we also output a majority *base-level* consensus sequence of all bases, not just SNPs, for each set of assigned reads by iterating through alignments in the BAM file. We return the ‘N’ base if the fraction of reads supporting the majority base is less than a parameter (default = 0.66).

### 2.5 Practical details and implementation

`devider` is implemented in Rust and uses the rust-htslib and rust-bio crates [39]. It is open-source and available at GitHub (https://github.com/bluenote-1577/devider) and through bioconda [40]. We implemented the following preset options to help users pick parameters: `old-long-reads, nanopore-r9, nanopore-r10, hi-fi`. These correspond to (*M, α, ρ*) = (10, 50, 0.02), (20, 150, 0.01), (35, 250, 0.005), and (100, 500, 0.001) respectively. The current default is `nanopore-r9`. We wrap `devider` in minimap2 and LoFreq in the `run_devider_pipeline` script in the repository. This runs minimap2, samtools, and LoFreq to generate an indexed BAM and VCF pair. For LoFreq, we found that disabling base-alignment qualities with the `-B` option improved sensitivity on nanopore reads. All other parameters were set to default.

## 3 Results

We show our benchmarking setups in **Supplementary Figure 3 and 4**. For a set of genomes, we used badread [41] to simulate nanopore long reads with default settings except for lengths and accuracy, which we make explicit for each dataset below. We picked an arbitrary reference genome and ran all tools with this reference genome. We then compared their predicted haplotypes to the true haplotypes. We benchmarked `devider` against iGDA v1.0.1, CliqueSNV v2.0.3, and RVHaplo v2. We tried running Strainline, a de novo viral quasispecies assembly method, but it outputted an error that was related to an unresolved GitHub issue (https://github.com/HaploKit/Strainline/issues/17). We ran all methods with default settings and iGDA in its Oxford Nanopore setting with the `ont_context_effect_read_qv_12_base_qv_12` model. Scripts for reproducing benchmarks and figures can be found at https://github.com/bluenote-1577/ devider-test. We ran all methods with 10 threads. We used the following metrics for benchmarking:

1. Hamming SNP error: the mean percentage of incorrect SNPs for each predicted haplotype against its best-matching genome.
2. Fraction recovered: the fraction of SNPs recovered for all genomes after matching predicted haplotypes against their best-matching genomes.
3. Earth mover’s distance (EMD) [42]: a measure of the distance between the predicted abundances and the true abundances. The pairwise distance function we used for the EMD is the number of mismatched SNPs between the predicted and true haplotype, and the weights are the predicted relative abundances.
4. Haplotype error: the predicted number of haplotypes minus the true number.

### 3.1 HIV-1 benchmarking (7 strains plus varying coverage and 2-30 strains)

HIV-1 serves as a standard benchmarking genome for viral quasispecies methods due to its fast mutation rate as an ssRNA virus and its clinical and public health importance. Thus, we created two HIV-1 communities: a 7-strain staggered abundance community and multiple communities from 2 to 30 strains with uniform abundances.

#### 7 strains at staggered abundances

We took a set of 30 HIV-1 genomes from Kinloch et al. [43] with accessions available in Supplementary Table 1. For the first 7 strain dataset, we selected an arbitrary reference (OR483991.1) and the 7 most similar strains as determined by skani [44]. The abundances were staggered at a 1:3:5:7:9:10:20 ratio, with the smallest strain coverage ranging from 3x to 160x. We simulated reads under three settings with (Accuracy, Length) = (95%, 9000bp), (98%, 9000bp), and (95%, 3000bp) with length standard deviation as 500bp. 95% accuracy represents older or faster nanopore sequencing runs, whereas 98% is more representative of the best current basecalling/chemistries [20]. HIV-1 genomes are approximately 9000 bp, so the two length settings represent complete and partial coverage.

On the 95% accuracy, 9000bp dataset (**Fig. 2 a**), `devider` had the best mean performance for fraction recovered (97.7% vs 86.2% for iGDA, the second best), the best haplotype error (−0.15 vs -0.92 for iGDA, the second best), and EMD (0.41 vs 5.89 for RVHaplo, the second best). iGDA and `devider` both achieved perfect a SNP Hamming error across all data points of 0%. The biggest performance difference was at low coverage; at 3x minimum coverage, `devider` could recover 6 haplotypes whereas all the other methods recovered 1, 1, and 3 haplotypes for RVHaplo, CliqueSNV, and iGDA respectively. We found that CliqueSNV consistently obtained 6 haplotypes, missing the low abundance haplotype. We tried to increase its sensitivity by lowering its abundance threshold to 0.25% (the same as `devider`), but it drastically overestimated the number of haplotypes, outputting > 30 haplotypes at high coverage. To show that `devider` is robust to parameter choices on this dataset, we varied the *k*-mer length between 10 to 30 and found that `devider` still had the best haplotype error, fraction recovered, and EMD (**Supplementary Fig. 5**).

**Fig. 2.**
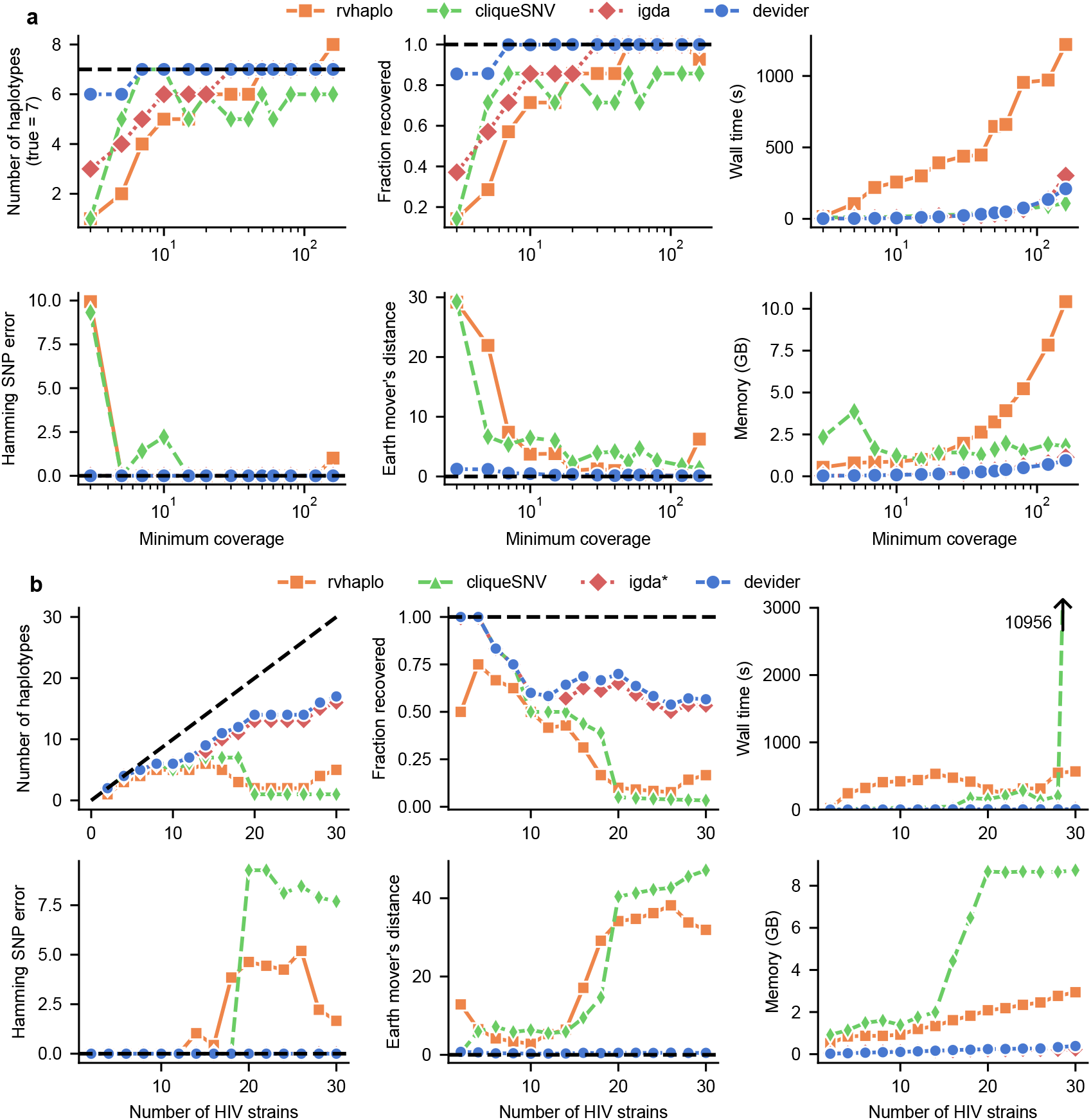
Benchmarking long-read haplotyping tools on simulated HIV-1 communities. **a**. 7 HIV-1 strains from Kinloch et al. [43] at staggered abundances (1:3:5:7:9:10:20) with simulated reads (9000 bp mean length; 95% mean accuracy). The x-axis indicates the depth of coverage for the lowest-coverage strain. **b**. 2 to 30 HIV-1 strains with 15x - 145x uniformly random coverage and the same simulated read lengths/accuracy. CliqueSNV failed to complete within the default 3-hour time limit for the last sample. *iGDA does not output abundances, so the earth mover’s distance could not be computed. Dashed black lines indicate optimal performance

On the 95%, 3000bp dataset (**Supplementary Fig. 6 a**), `devider` was the best for mean fraction recovered (92.3% vs 83.5% for the second-best RVHaplo) and EMD (1.18 vs 5.28 for the second-best RVHaplo). `devider` had the smallest absolute mean haplotype error (+0.15) compared to the other methods; other methods all underestimated the number of haplotypes (iGDA was the second best at -0.46). However, the mean Hamming SNP error for `devider` (0.37%) was slightly worse than iGDA (0.06%) and RVHaplo (0.07%).

On the 98% accuracy, 9000bp dataset (**Supplementary Fig. 6 b**), `devider` was the best method on every metric with iGDA in second. `devider` had near-perfect performance in terms of mean fraction recovered (98.9% vs 88.1% for iGDA), haplotype error (−0.08 vs -0.69 for iGDA), Hamming SNP error (0% vs 0.024% for iGDA) and EMD (0.31 vs 4.4 for CliqueSNV, the second best). de Bruijn graph methods work well with lower error rates because the probability of having an error within a k-mer is approximately (*accuracy*)^*k*^; this is encouraging for future nanopore datasets as read accuracy increases.

#### 2-30 strains at uniform coverages

We simulated reads at 95% accuracy and 9000 bp average length for 2-30 strains from the same set of HIV-1 genomes, with each genome at 15x-145x coverage uniformly at random (**Fig. 2 b**). `devider` was the best method on all mean metrics for this dataset with iGDA a close second. CliqueSNV also performed well when the number of strains was < 10 (81.7% mean fraction recovered compared to 83.6% and 83.4% for `devider` and iGDA), but its performance dropped when more than 10 strains were present (37.5% mean fraction recovered for 10-20 strains versus 65.6% for `devider`).

`devider` and iGDA also stood out in terms of efficiency compared to the other methods. iGDA and `devider` took 5 and 23 seconds on average respectively and < 0.5 GB of RAM. Note that we included lofreq’s runtime and memory usage in `devider`’s results. RVHaplo took 370 seconds and CliqueSNV 820 seconds on average. CliqueSNV was efficient except for the case with 30 strains, where the runtime ballooned to 10956 seconds. We found that this was because CliqueSNV sets a 3 hour (10800 seconds) time limit by default if it can not solve the haplotyping problem, after which it outputs no haplotypes.

#### 3.2 Synthetic AMR gene haplotyping for 53 AMR gene groups

We chose AMR genes as a gene-level haplotyping benchmark due to their diversity in sequence composition, length, biological significance, and also the prevalence of targeted enrichment sequencing protocols [45,46,47] for which our methods are potentially usable. We clustered AMR genes in the MEGARes database version 3.0 [48] by computing all pairwise average nucleotide identities (ANI) and mean alignment fractions (AF) with skani (using the `slow` preset) and then using the Leiden algorithm [49] with edge weights as *ANI* \**AF* at resolution 1.00. Of the remaining 53 clusters with ≥15 different haplotypes, we sampled between 2-15 haplotypes at coverage 80x-1000x, both uniformly at random. We then simulated reads at 95% accuracy and 1500 bp length with 200bp standard deviation. The AMR gene lengths ranged from 721 - 3303 bp.

On this dataset, `devider` was the best method for mean haplotype error, SNP Hamming error, and fraction recovered (**Fig. 3 a**), with ‒1.5, 0.06% and 83.3% respectively. CliqueSNV was the next best method with ‒4.2, 0.34%, and 60.2% on the same metrics. To investigate the discrepancy between different methods, we stratified the fraction recovered by coverage, % identity of haplotype to reference, and abundance (**Fig. 3 b**). As expected, lower coverage, which is correlated with lower abundance, leads to lower fraction recovered for all methods. We found the likely cause of the performance discrepancy to be high-similarity haplotypes: > 2/3 of the haplotypes had > 99.5% similarity to the reference, and on these haplotypes, `devider`’s fraction recovered was 73.8% compared to CliqueSNV with 46.3%, the next best method. Thus, `devider` is the most sensitive method for highly similar haplotypes, which is important because capturing even small variations in AMR genes can alter phenotypes [4].

**Fig. 3.**
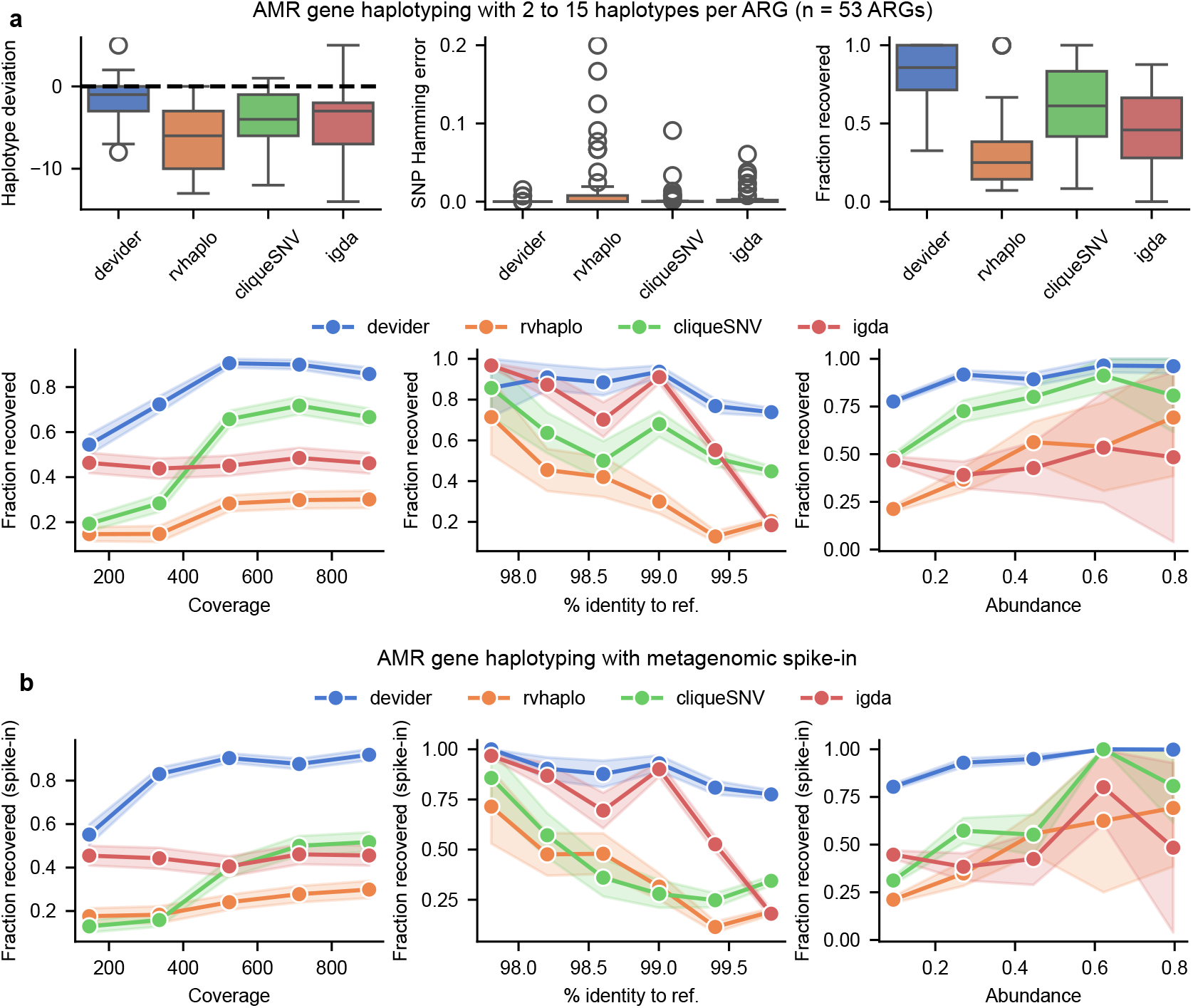
Benchmarking for haplotyping of synthetic mixtures of antimicrobial resistance (AMR) genes. **a**. Top: results for 53 sets of AMR genes with 2-15 haplotypes and reads simulated at 80x - 1000x coverage (95% identity; 1500 mean length), both picked uniformly at random. Bottom: the same results but with fraction recovered as a function of the haplotype’s coverage, its % nucleotide identity to the reference, and its abundance (i.e., normalized coverage). **b**. Fraction recovered for the AMR genes after spiking the AMR reads into a synthetic long-read mouse gut metagenome from CAMI2. Each method was rerun after aligning the pooled dataset against the AMR gene references. Error bars indicate standard errors after binning data points along the x-axis. Box plots show the median, the 25th and 75th percentiles, and 1.5× the interquartile range.

#### AMR dataset with spike-in metagenome

We extend the previous AMR haplotyping experiment to a metagenomics setting. In metagenomics, the input is a mixture of microbial genomic reads *and* reads from AMR genes. Thus, we mixed the previously simulated AMR reads into a simulated long-read mouse gut metagenome from CAMI2 [50] (labeled as sample 0), hereafter referred to as the spike-in metagenome.

Notably, genomes within the CAMI2 metagenome contain AMR genes or sequences with homology to AMR genes. Mapping the CAMI2 reads against MEGARES with minimap2 resulted in 130 AMR genes with >2x coverage but only 10 AMR genes with > 20x coverage. Thus, the spike-in metagenome contains the previously simulated AMR haplotypes (with 2-15 haplotypes and 80-1000x coverage) as well as this new tail of low-abundance AMR haplotypes from the CAMI2 metagenome.

Under this setup (shown in **Supplementary Fig. 4**), we measured each method’s ability to recover the original simulated AMR haplotypes (**Fig. 3 b**). `devider`, iGDA, and RVHaplo had < 2% difference in fraction recovered compared to the previous (without spike-in) case. However, CliqueSNV fell to 42.9% (with spike-in) from 60.2% (no spike-in). Thus, `devider` can recover abundant haplotypes for a long-tailed abundance distribution with low abundance haplotypes, a common characteristic of metagenomics data.

### 3.3 Results on real heterogeneous sequencing mixtures

#### HIV-1 co-infection haplotyping

Mori et al. [51] sequenced a set of HIV-1 samples with full-length nanopore amplicons (5.8-7% sequencing error rate) and detected a potential HIV-1 co-infection for a patient (labeled TRN9) by stochastically subsampling reads and reassembling. Here, we ran `devider`, iGDA, RVHaplo, and CliqueSNV on these reads and an arbitrary reference (NC_001802.1) to try to confirm their results. `devider` gave four haplotypes (60.0%, 32.34%, 4.12%, and 3.57% abundance) and RVHaplo gave two (67% and 32.9% abundance). CliqueSNV only output 1 haplotype and iGDA output no haplotypes.

Mori et al. found two majority haplotypes, which is corroborated by RVHaplo and `devider`. We show `devider`’s two majority haplotypes in **Fig. 4 a**. We investigated the two additional minority haplotypes output by `devider` to see if they were erroneous; curiously, these two haplotypes were a mix of the two majority haplotypes and well supported by “chimeric” reads (**Supplementary Fig. 7**). HIV-1 is known to recombine, which may be a possible explanation, but we do not speculate further since we can not rule out sequencing artifacts.

**Fig. 4.**
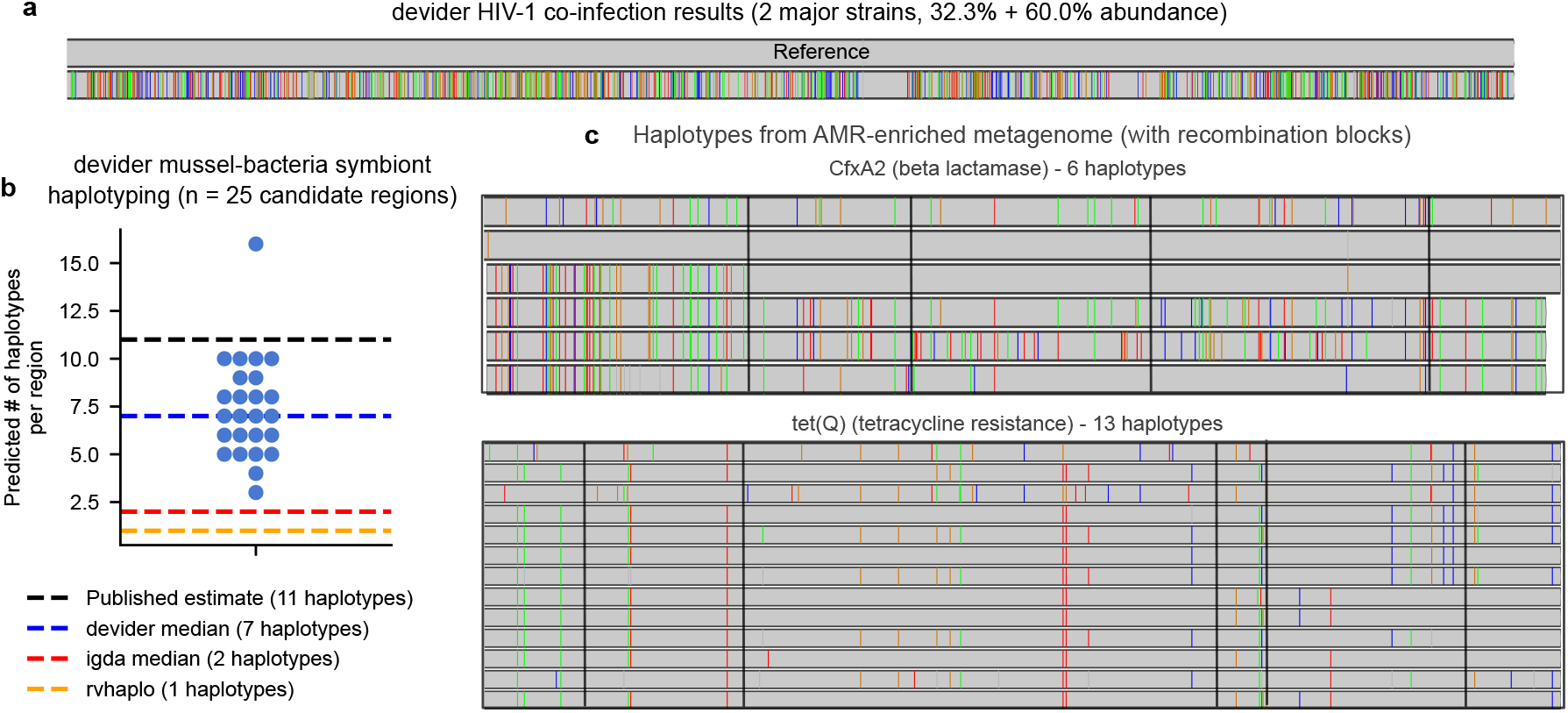
Long-read haplotyping results from real samples subjected to a variety of sequencing technologies. **a**. Haplotypes from long-read HIV-1 nanopore sequencing (93 - 94.2% predicted sequencing accuracy) of an HIV co-infection from Mori et al. [51]. Two major haplotypes were found by `devider`, confirming previous results. Mismatched bases are shown with the reference as the upper haplotype. **b**. Haplotyping results for PacBio RS II sequencing (89.5% mean gap-compressed identity against reference) of an intracellular bacterial symbiont community within deep-sea mussels from Ansorge et al. [52], who predicted 11 strains to be present. 25 candidate single-copy regions with high SNP diversity were haplotyped by `devider`, iGDA, and RVHaplo; CliqueSNV was excluded because it timed out on multiple regions. `devider` produced higher diversity estimates than iGDA and RVHaplo, which both produced ≤3 haplotypes across all sites. **c**. `devider` haplotyping of a long-read bovine gut metagenome enriched for AMR genes. CfxA2 (3200x coverage) and tet(Q) (19500x coverage and last 1000bp shown) haplotype sequences with > 30x coverage and 1% abundance are shown with mismatches against their reference sequences in MEGARES v3.0. Mismatches shared by all haplotypes are removed. Recombination blocks are outlined in black as predicted by GARD.

#### Mussel-bacteria symbiont community with 11 estimated strains

Ansorge et al. [52] investigated the strain-level diversity of an intracellular, sulfur-oxidizing bacterial symbiont community within deep-sea mussels. They sequenced one sample using the PacBio RS II, which produces high-error long reads. On this sample, they estimated 11 strains by using an assembly approach and then counting single-copy genes. We investigated if haplotyping could give similar estimates, even with noisy reads (mean gap-compressed identity against reference = 89.5%). To generate reasonable diversity estimates, we haplotyped 3kb regions of the reference genome (GCF_900128535) that had > 10 SNPs (as detected by LoFreq) and coverage between 200 and 250. These criteria were chosen because the average read length was 3.8kb and the estimated single-copy coverage, calculated by dividing the total read bases by the genome size, was 220. We ran all methods on these regions in PacBio mode (if such a preset existed) and the `old-long-reads` preset for `devider`.

On these 25 regions, `devider` estimated a median of seven haplotypes (**Fig. 4 b**), whereas iGDA and RVHaplo estimated ≤ 2 median haplotypes. CliqueSNV failed to run on multiple regions, timing out after 3 hours and outputting nothing, so we did not include its results. `devider` estimated 16 haplotypes for one region, and subsequent investigation revealed many supplementary alignments near the edge of contigs. The first 600 bases had mean coverage > 300x, possibly indicating a duplicated region (**Supplementary Fig. 8**) for a subset of the strains. Ultimately, `devider` was able to capture some of the known diversity in these samples using noisy long reads, whereas other methods failed.

#### Discovering recombinant AMR genes in AMR-enriched long-read metagenomes

We used `devider` to haplotype an AMR-enriched PacBio CCS long-read fecal metagenome from a cow that received intensive antibiotic treatments [45]. We first dereplicated the MEGARes v3.0 database at 95% using vsearch [53] to avoid ambiguous mapping of reads to highly similar genes. We then ran `devider` with extra stringent parameters, setting the minimum abundance to 1% and minimum depth to 30x.

In total, we found 18 different dereplicated AMR genes with ≥ 2 haplotypes. Of these 18 genes, 9 were tetracycline resistance genes that were phased into 52 distinct haplotypes. The highest coverage gene was tet(Q) [54] at 19500x coverage and phased into 13 haplotypes. Other high-diversity genes we found included mefA [55], a macrolide efflux pump, at 7100x coverage and phased into 12 haplotypes; as well as CfxA2 [56], a beta-lactamase, at 3200x coverage and phased into 6 haplotypes. We illustrate the haplotypes of tet(Q) and CfxA2 in **Fig. 4 c** (IGV [57] screenshots in **Supplementary Fig. 9 and 10**). We found a distinct mosaic structure within these haplotypes, suggesting a history of recombination within these haplotypes. We used MAFFT [58] to generate a multiple sequence alignment from `devider`’s haplotypes and GARD [59] to detect recombination, which found evidence of recombination for both genes. We draw breakpoints where GARD’s model-averaged support was > 0.3 in **Fig. 4 c**. Mosaism due to recombination is a well-documented characteristic of some ribosomal protection proteins including tet(Q) [60], and CfxA genes are commonly colocalized with an element known to play a role in the mosaic behavior of conjugative elements [61]. Thus, these detected recombination events are supported by known mechanisms in these two AMR genes.

## 4 Conclusion

We presented `devider`, a method for retrieving high-similarity haplotypes from long-read sequencing of heterogeneous sequences. `devider` leverages a positional de Bruijn graph (PDBG) assembly approach on a subset of informative alleles to disentangle variation. This framework is efficient and naturally resolves variation without the need to explicitly infer the number of haplotypes. The key technical challenge was to remove sequencing artifacts within the PDBG, especially for error-prone long reads, while retaining high sensitivity, which we accomplished through an error-aware sequence-to-graph alignment approach.

We designed `devider` to work with a wider range of technologies and sequencing error rates. We limited `devider` to reconstructing “small” sequences on the order of read length for conservative recovery. As error rates improve, it may be possible to attempt longer haplotype reconstruction using our approach. Another limitation is our reference-based approach, which is not able to recover new sequences de novo. Nevertheless, reference-based approaches are intrinsically more efficient and simpler than de novo approaches, and we believe they are complementary to de novo approaches. As the capabilities and the need for haplotype-level resolved sequences from long reads continue to increase, `devider` will be a fast, useful tool for retrieving accurate haplotypes.

## References

1. Domingo, E. & Perales, C. Viral quasispecies. PLoS Genetics 15, e1008271 (2019).

2. Cuevas, J. M., Geller, R., Garijo, R., López-Aldeguer, J. & Sanjuán, R. Extremely High Mutation Rate of HIV-1 In Vivo. PLoS Biology 13, e1002251 (2015).

3. Van Rossum, T., Ferretti, P., Maistrenko, O. M. & Bork, P. Diversity within species: Interpreting strains in microbiomes. Nature Reviews Microbiology 18, 491–506 (2020).

4. Vedantam, G., Guay, G. G., Austria, N. E., Doktor, S. Z. & Nichols, B. P. Characterization of mutations contributing to sulfathiazole resistance in Escherichia coli. Antimicrobial Agents and Chemotherapy 42, 88–93 (1998).

5. Olkkola, S., Juntunen, P., Heiska, H., Hyytiäinen, H. & Hänninen, M.-L. Mutations in the rpsL gene are involved in streptomycin resistance in Campylobacter coli. Microbial Drug Resistance (Larchmont, N.Y.) 16, 105–110 (2010).

6. Bickhart, D. M. et al. Generating lineage-resolved, complete metagenome-assembled genomes from complex microbial communities. Nature Biotechnology 40, 711–719 (2022).

7. Cheng, H., Concepcion, G. T., Feng, X., Zhang, H. & Li, H. Haplotype-resolved de novo assembly using phased assembly graphs with hifiasm. Nature Methods 18, 170–175 (2021).

8. Feng, X., Cheng, H., Portik, D. & Li, H. Metagenome assembly of high-fidelity long reads with hifiasm-meta. Nature Methods 19, 671–674 (2022).

9. Benoit, G. et al. High-quality metagenome assembly from long accurate reads with metaMDBG. Nature Biotechnology 1–6 (2024).

10. Li, H. & Durbin, R. Genome assembly in the telomere-to-telomere era. Nature Reviews Genetics 25, 658–670 (2024).

11. Feng, Z., Clemente, J. C., Wong, B. & Schadt, E. E. Detecting and phasing minor single-nucleotide variants from long-read sequencing data. Nature Communications 12, 3032 (2021).

12. Knyazev, S. et al. Accurate assembly of minority viral haplotypes from next-generation sequencing through efficient noise reduction. Nucleic Acids Research 49, e102 (2021).

13. Cai, D. & Sun, Y. Reconstructing viral haplotypes using long reads. Bioinformatics 38, 2127–2134 (2022).

14. Lin, J.-H., Chen, L.-C., Yu, S.-C. & Huang, Y.-T. LongPhase: An ultra-fast chromosome-scale phasing algorithm for small and large variants. Bioinformatics 38, 1816–1822 (2022).

15. Edge, P., Bafna, V. & Bansal, V. HapCUT2: Robust and accurate haplotype assembly for diverse sequencing technologies. Genome Research 27, 801–812 (2017).

16. Shaw, J. & Yu, Y. W. Flopp: Extremely Fast Long-Read Polyploid Haplotype Phasing by Uniform Tree Partitioning. Journal of Computational Biology 29, 195–211 (2022).

17. Lancia, G., Bafna, V., Istrail, S., Lippert, R. & Schwartz, R. SNPs Problems, Complexity, and Algorithms. In auf der Heide, F. M. (ed.) Algorithms — ESA 2001, Lecture Notes in Computer Science, 182–193 (Springer, Berlin, Heidelberg, 2001).

18. Zhou, Q. et al. KSNP: A fast de Bruijn graph-based haplotyping tool approaching data-in time cost. Nature Communications 15, 3126 (2024).

19. Patterson, M. et al. WhatsHap: Weighted Haplotype Assembly for Future-Generation Sequencing Reads. Journal of Computational Biology: A Journal of Computational Molecular Cell Biology 22, 498–509 (2015).

20. Sereika, M. et al. Oxford Nanopore R10.4 long-read sequencing enables the generation of near-finished bacterial genomes from pure cultures and metagenomes without short-read or reference polishing. Nature Methods 19, 823–826 (2022).

21. Zagordi, O., Bhattacharya, A., Eriksson, N. & Beerenwinkel, N. ShoRAH: Estimating the genetic diversity of a mixed sample from next-generation sequencing data. BMC Bioinformatics 12, 119 (2011).

22. Eliseev, A. et al. Evaluation of haplotype callers for next-generation sequencing of viruses. Infection, genetics and evolution : journal of molecular epidemiology and evolutionary genetics in infectious diseases 82, 104277 (2020).

23. Shaw, J., Gounot, J.-S., Chen, H., Nagarajan, N. & Yu, Y. W. Floria: Fast and accurate strain haplotyping in metagenomes. Bioinformatics 40, i30–i38 (2024).

24. Vicedomini, R., Quince, C., Darling, A. E. & Chikhi, R. Strainberry: Automated strain separation in low-complexity metagenomes using long reads. Nature Communications 12, 4485 (2021).

25. Kazantseva, E., Donmez, A., Frolova, M., Pop, M. & Kolmogorov, M. Strainy: Phasing and assembly of strain haplotypes from long-read metagenome sequencing. Nature Methods 1–10 (2024).

26. Li, H. Minimap2: Pairwise alignment for nucleotide sequences. Bioinformatics 34, 3094–3100 (2018).

27. Wilm, A. et al. LoFreq: A sequence-quality aware, ultra-sensitive variant caller for uncovering cell-population heterogeneity from high-throughput sequencing datasets. Nucleic Acids Research 40, 11189–11201 (2012).

28. Guo, Y. et al. The effect of strand bias in Illumina short-read sequencing data. BMC Genomics 13, 666 (2012).

29. Benjamini, Y. & Hochberg, Y. Controlling the False Discovery Rate: A Practical and Powerful Approach to Multiple Testing. Journal of the Royal Statistical Society. Series B (Methodological) 57, 289–300 (1995). 2346101.

30. McElroy, K., Zagordi, O., Bull, R., Luciani, F. & Beerenwinkel, N. Accurate single nucleotide variant detection in viral populations by combining probabilistic clustering with a statistical test of strand bias. BMC Genomics 14, 501 (2013).

31. Liu, D. & Steinegger, M. Block aligner: Fast and flexible pairwise sequence alignment with SIMD-accelerated adaptive blocks. Preprint, Bioinformatics (2021).

32. Ronen, R., Boucher, C., Chitsaz, H. & Pevzner, P. SEQuel: Improving the accuracy of genome assemblies. Bioinformatics 28, i188–i196 (2012).

33. Bao, E., Jiang, T. & Girke, T. AlignGraph: Algorithm for secondary de novo genome assembly guided by closely related references. Bioinformatics 30, i319–i328 (2014).

34. Cameron, D. L. et al. GRIDSS: Sensitive and specific genomic rearrangement detection using positional de Bruijn graph assembly. Genome Research 27, 2050 (2017).

35. Myers, E. W. The fragment assembly string graph. Bioinformatics (Oxford, England) 21 Suppl 2, ii79–85 (2005).

36. Li, D., Liu, C.-M., Luo, R., Sadakane, K. & Lam, T.-W. MEGAHIT: An ultra-fast single-node solution for large and complex metagenomics assembly via succinct de Bruijn graph. Bioinformatics 31, 1674–1676 (2015).

37. Stevenson, K. R., Coolon, J. D. & Wittkopp, P. J. Sources of bias in measures of allele-specific expression derived from RNA-seq data aligned to a single reference genome. BMC Genomics 14, 536 (2013).

38. Delahaye, C. & Nicolas, J. Sequencing DNA with nanopores: Troubles and biases. PLOS ONE 16, e0257521 (2021).

39. Köster, J. Rust-Bio: A fast and safe bioinformatics library. Bioinformatics 32, 444–446 (2016).

40. Grüning, B. et al. Bioconda: Sustainable and comprehensive software distribution for the life sciences. Nature Methods 15, 475–476 (2018).

41. Wick, R. R. Badread: Simulation of error-prone long reads. Journal of Open Source Software 4, 1316 (2019).

42. Rubner, Y., Tomasi, C. & Guibas, L. A metric for distributions with applications to image databases. In Sixth International Conference on Computer Vision (IEEE Cat. No.98CH36271), 59–66 (1998).

43. Kinloch, N. N. et al. HIV reservoirs are dominated by genetically younger and clonally enriched proviruses. mBio 14, e02417–23 (2023).

44. Shaw, J. & Yu, Y. W. Fast and robust metagenomic sequence comparison through sparse chaining with skani. Nature Methods 1–5 (2023).

45. Slizovskiy, I. B. et al. Target-enriched long-read sequencing (TELSeq) contextualizes antimicrobial resistance genes in metagenomes. Microbiome 10, 185 (2022).

46. Shay, J. A. et al. Exploiting a targeted resistome sequencing approach in assessing antimicrobial resistance in retail foods. Environmental Microbiome 18, 25 (2023).

47. Baba, H., Kuroda, M., Sekizuka, T. & Kanamori, H. Highly sensitive detection of antimicrobial resistance genes in hospital wastewater using the multiplex hybrid capture target enrichment. mSphere 8, e00100–23 (2023).

48. Bonin, N. et al. MEGARes and AMR++, v3.0: An updated comprehensive database of antimicrobial resistance determinants and an improved software pipeline for classification using high-throughput sequencing. Nucleic Acids Research 51, D744–D752 (2023).

49. Traag, V. A., Waltman, L. & van Eck, N. J. From Louvain to Leiden: Guaranteeing well-connected communities. Scientific Reports 9, 5233 (2019).

50. Meyer, F. et al. Critical Assessment of Metagenome Interpretation: The second round of challenges. Nature Methods 19, 429–440 (2022).

51. Mori, M. et al. Nanopore Sequencing for Characterization of HIV-1 Recombinant Forms. Microbiology Spectrum 10, e01507–22 (2022).

52. Ansorge, R. et al. Functional diversity enables multiple symbiont strains to coexist in deep-sea mussels. Nature Microbiology 4, 2487–2497 (2019).

53. Rognes, T., Flouri, T., Nichols, B., Quince, C. & Mahé, F. VSEARCH: A versatile open source tool for metagenomics. PeerJ 4, e2584 (2016).

54. Lacroix, J.-M. & Walker, C. B. Detection and prevalence of the tetracycline resistance determinant Tet Q in the microbiota associated with adult periodontitis. Oral Microbiology and Immunology 11, 282–288 (1996).

55. Daly, M. M., Doktor, S., Flamm, R. & Shortridge, D. Characterization and Prevalence of MefA, MefE, and the Associated msr(D) Gene in Streptococcus pneumoniae Clinical Isolates. Journal of Clinical Microbiology 42, 3570–3574 (2004).

56. Iwahara, K. et al. Detection of cfxA and cfxA2, the β-Lactamase Genes of Prevotella spp., in Clinical Samples from Dentoalveolar Infection by Real-Time PCR. Journal of Clinical Microbiology 44, 172–176 (2006).

57. Robinson, J. T. et al. Integrative Genomics Viewer. Nature biotechnology 29, 24–26 (2011).

58. Katoh, K., Misawa, K., Kuma, K.-i. & Miyata, T. MAFFT: A novel method for rapid multiple sequence alignment based on fast Fourier transform. Nucleic Acids Research 30, 3059–3066 (2002).

59. Kosakovsky Pond, S. L., Posada, D., Gravenor, M. B., Woelk, C. H. & Frost, S. D. GARD: A genetic algorithm for recombination detection. Bioinformatics 22, 3096–3098 (2006).

60. Warburton, P. J., Amodeo, N. & Roberts, A. P. Mosaic tetracycline resistance genes encoding ribosomal protection proteins. Journal of Antimicrobial Chemotherapy 71, 3333–3339 (2016).

61. García, N. et al. Genetic determinants for cfxA expression in Bacteroides strains isolated from human infections. Journal of Antimicrobial Chemotherapy 62, 942–947 (2008).

