## Supplementary Materials for "devider: long-read reconstruction of many diverse haplotypes"

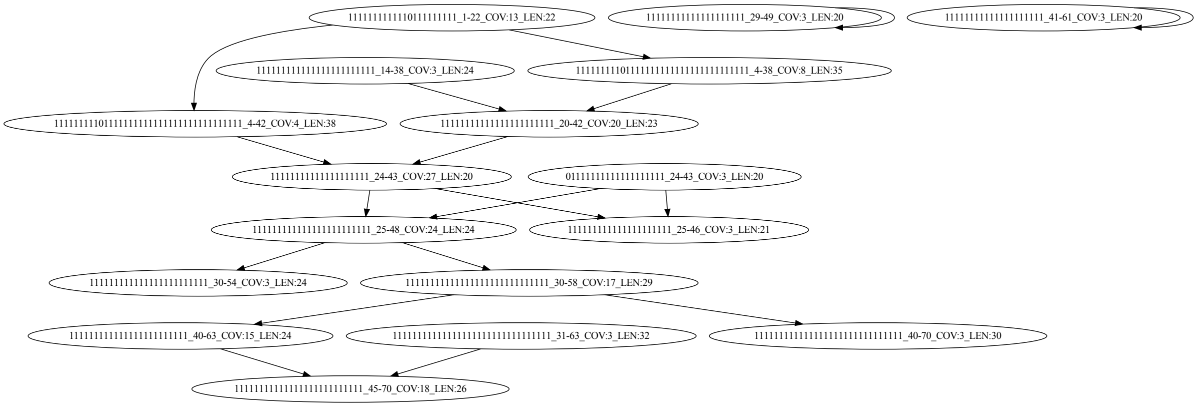

**Supplementary Figure 1.** Unitig graph for long reads (95% accuracy) for a single HIV-1 genome. The k-mer coverage cutoff was 2. The format is (alleles)_(start-end)_COV:(coverage)_LEN:(number of alleles spanned). Notice that low coverage nodes and nodes with deletions (i.e., end - start + 1 $\neq$ length) are more likely to be erroneous.

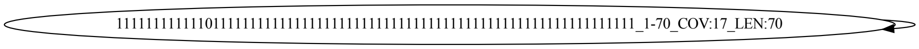

**Supplementary Figure 2.** Cleaned unitig graph for long reads (95% accuracy) for a single HIV-1 genome. The graph contains one unitig, as expected. The format is (alleles)_(start-end)_COV:(coverage)_LEN:(number of alleles spanned).

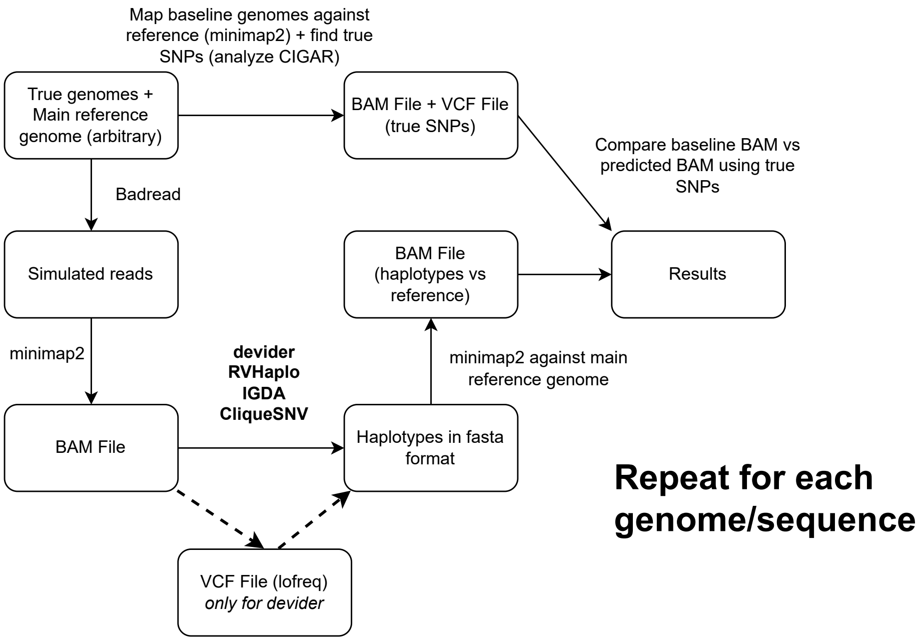


**Supplementary Figure 3.** Benchmarking flowchart for the HIV-1 and antimicrobial resistance gene synthetic benchmarks.

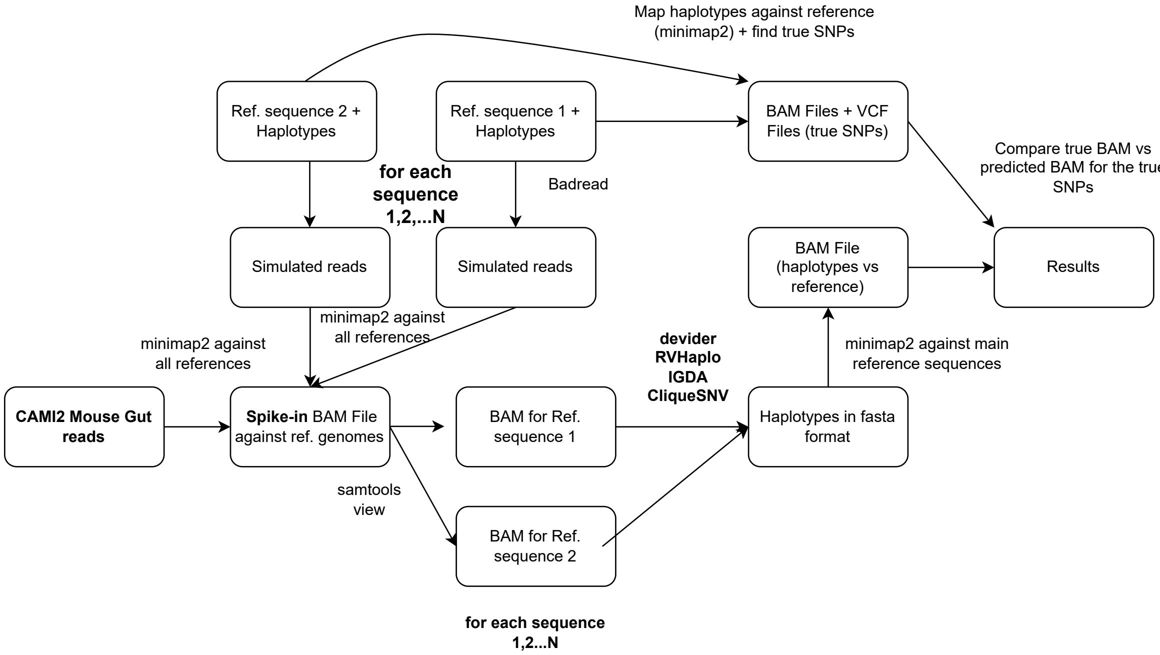


**Supplementary Figure 4.** Benchmarking flowchart for the antimicrobial resistance gene **spike-in** synthetic benchmarks. The AMR gene reads were spiked into a mouse gut metagenome, and BAM files were generated for each AMR gene of interest with `samtools view` prior to haplotyping.

| Genome accessions for HIV communities |
|---|
| - OR483999 |
| - OR483988 |
| - OR483990 |
| - OR483987 |
| - OR484006 |
| - OR484007 |
| - OR484012 |
| - OR484013 |
| - OR484014 |
| - OR484015 |
| - OR484016 |
| - OR483993 |
| - OR483994 |
| - OR483989 |
| - OR483992 |
| - OR483997 |
| - OR484000 |
| - OR484002 |
| - OR483986 |
| - OR484017 |
| - OR484018 |
| - OR484019 |
| - OR484020 |
| - OR484021 |
| - OR484022 |
| - OR484001 |
| - OR483985 |
| - OR484004 |
| - OR484005 |
| - OR483996 |

**Supplementary Table 1.** Genomes used for the HIV community. The first 7 (delimited by an horizontal line) are used for the 7 strain community and all 30 are used when benchmarking communities with 2 - 30 strains.

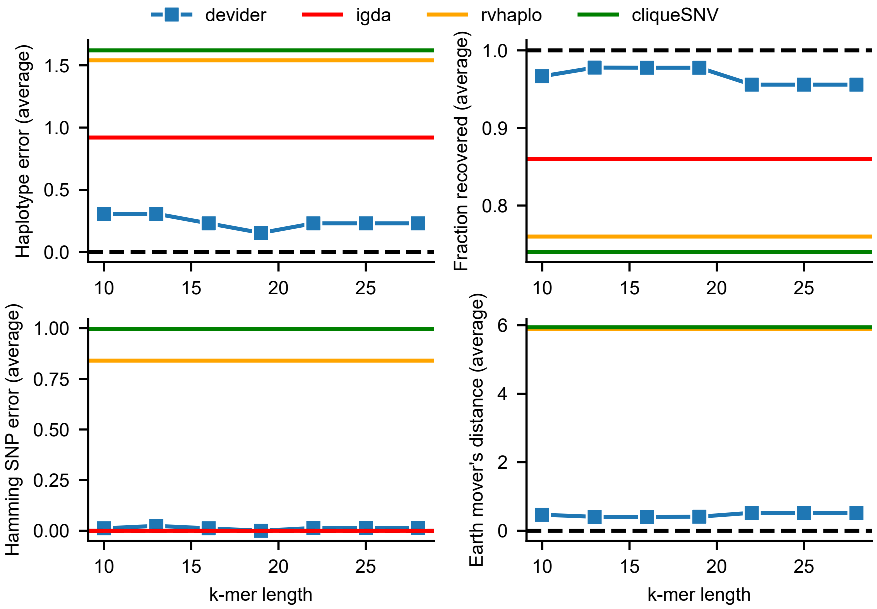


**Supplementary Figure 5.** devider's results over varying values of $k$-mer length $k$ on the 7-strain HIV community with 9000 bp length and 95% accuracy simulated reads. devider dynamically selects $k$ based on the data by default; instead, we fixed $k$ from 10 to 30 and show averaged results over all data points. The other three methods are displayed as constant horizontal lines. The dashed black line shows the optimal values.

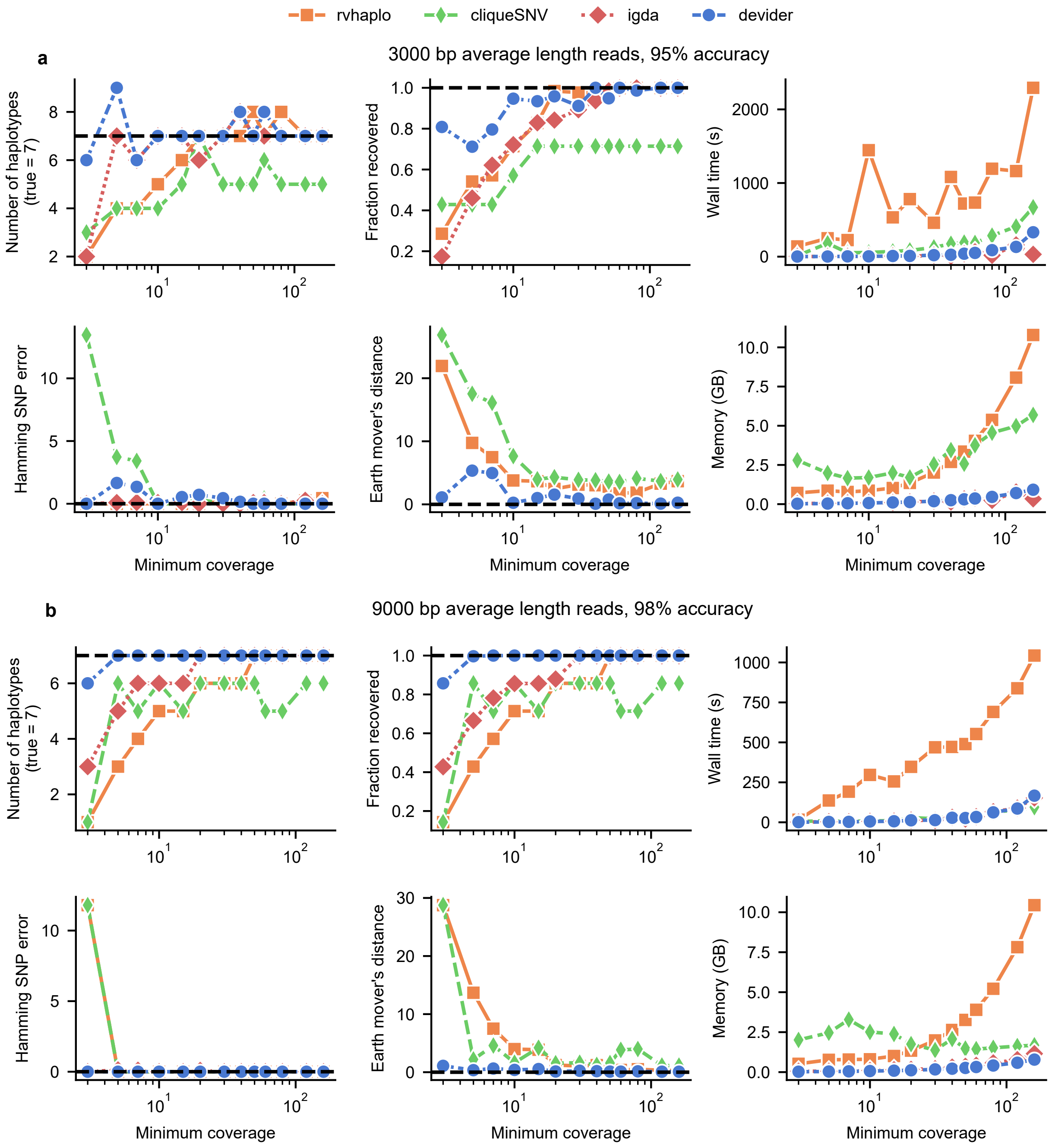


**Supplementary Figure 6.** The same experiment as Fig. 2a in the main paper on the 7-strain HIV synthetic community with staggered abundances. **a.** simulated reads have 3000bp average length and 95% accuracy. **b.** simulated reads have 9000 bp average length and 98% accuracy.

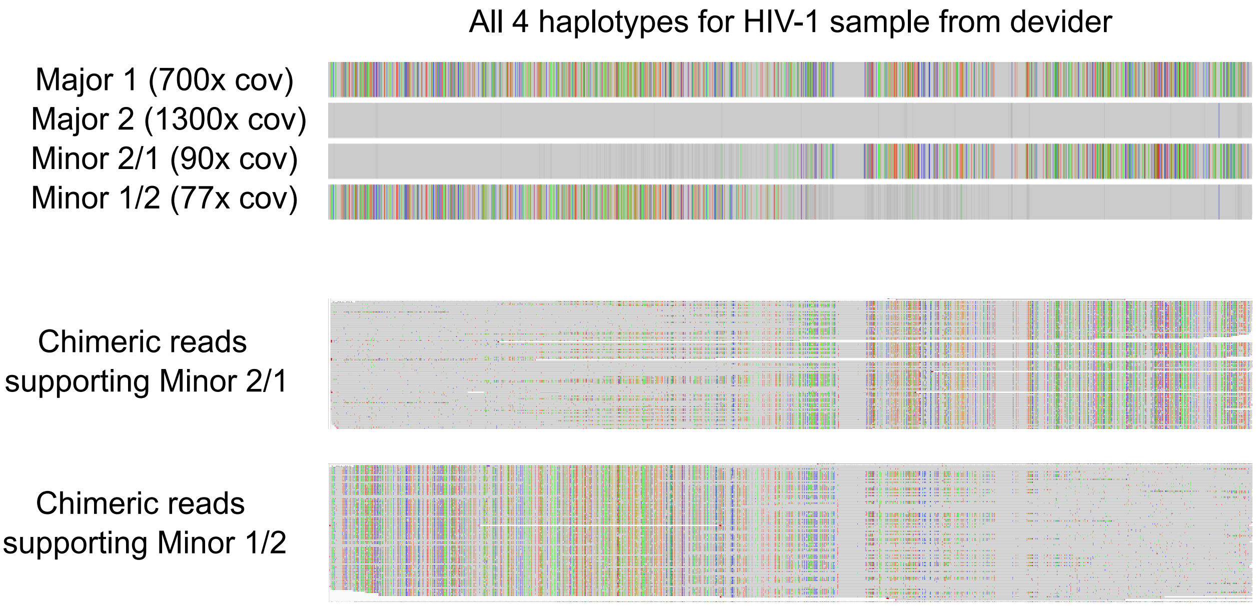


**Supplementary Figure 7.** All four haplotypes found by devider on the HIV-1 co-infection dataset. Two major haplotypes were found and two minor haplotypes were also found. Gray bars indicate alleles with $< 66\%$ of the reads supporting that allele. For the "chimeric" minor haplotypes, supporting reads assigned by devider are shown. The reference sequence for all alignments is a consensus generated by abPOA [1] from Major Haplotype 2's reads.

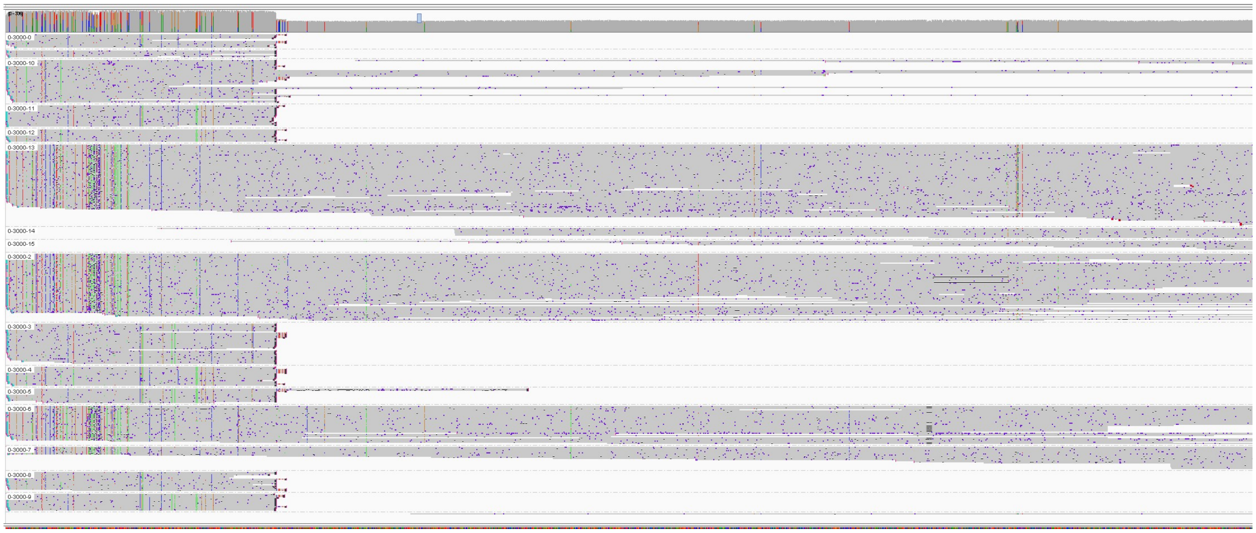

**Supplementary Figure 8.** A region with 16 estimated haplotypes by devider for the mussel-bacterial symbiont community. An elevated region of high coverage at the start of the contig (mean $>$ 300x coverage vs estimated 220x single-copy coverage) and supplementary mapping reads suggest a duplicated region, which makes the number of haplotypes estimated higher.

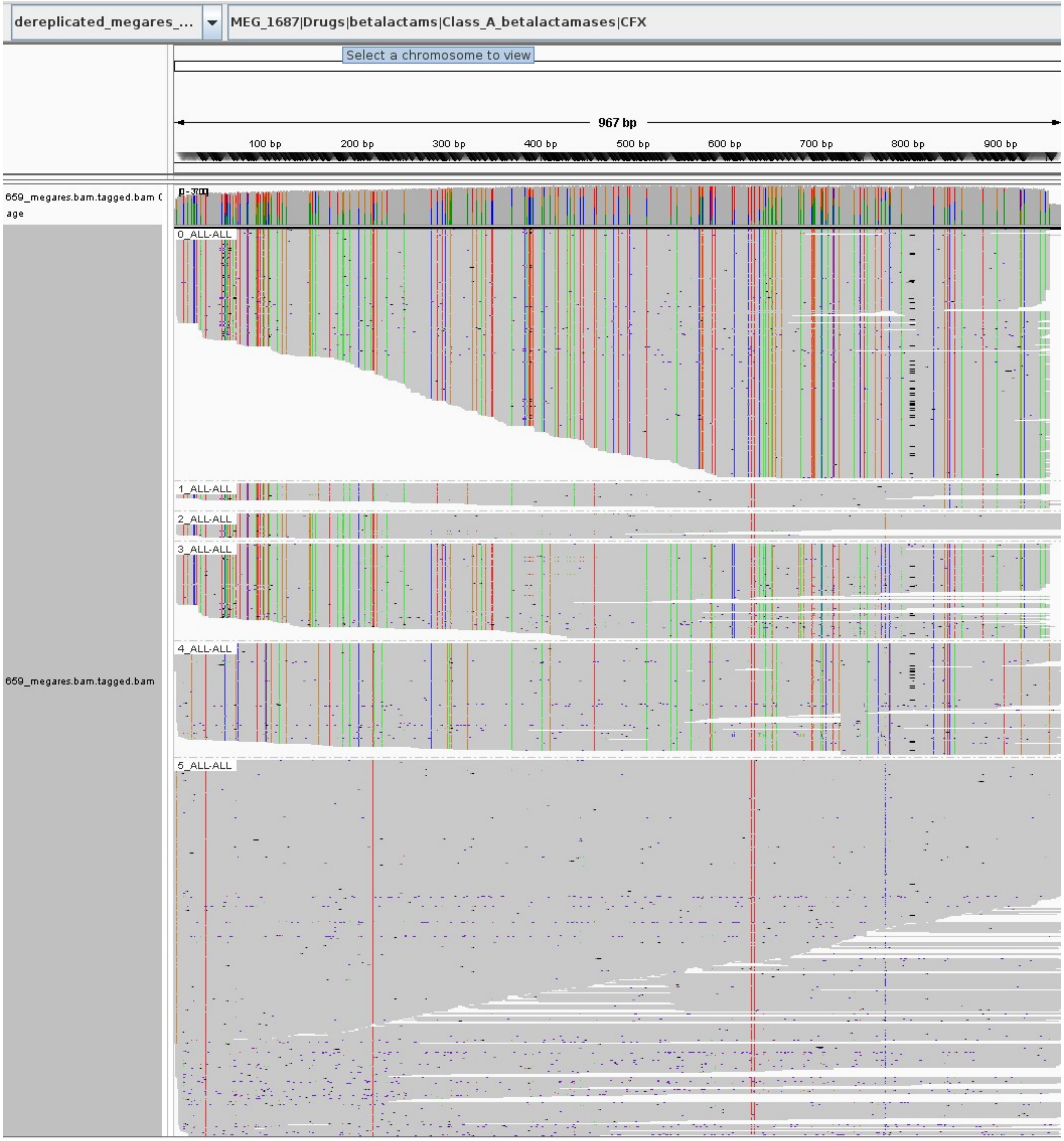


**Supplementary Figure 9.** Reads supporting the CfxA2 haplotypes shown in Fig. 4 c. Reads were downsampled to 1000x coverage per 1000 bp window for visualization purposes.

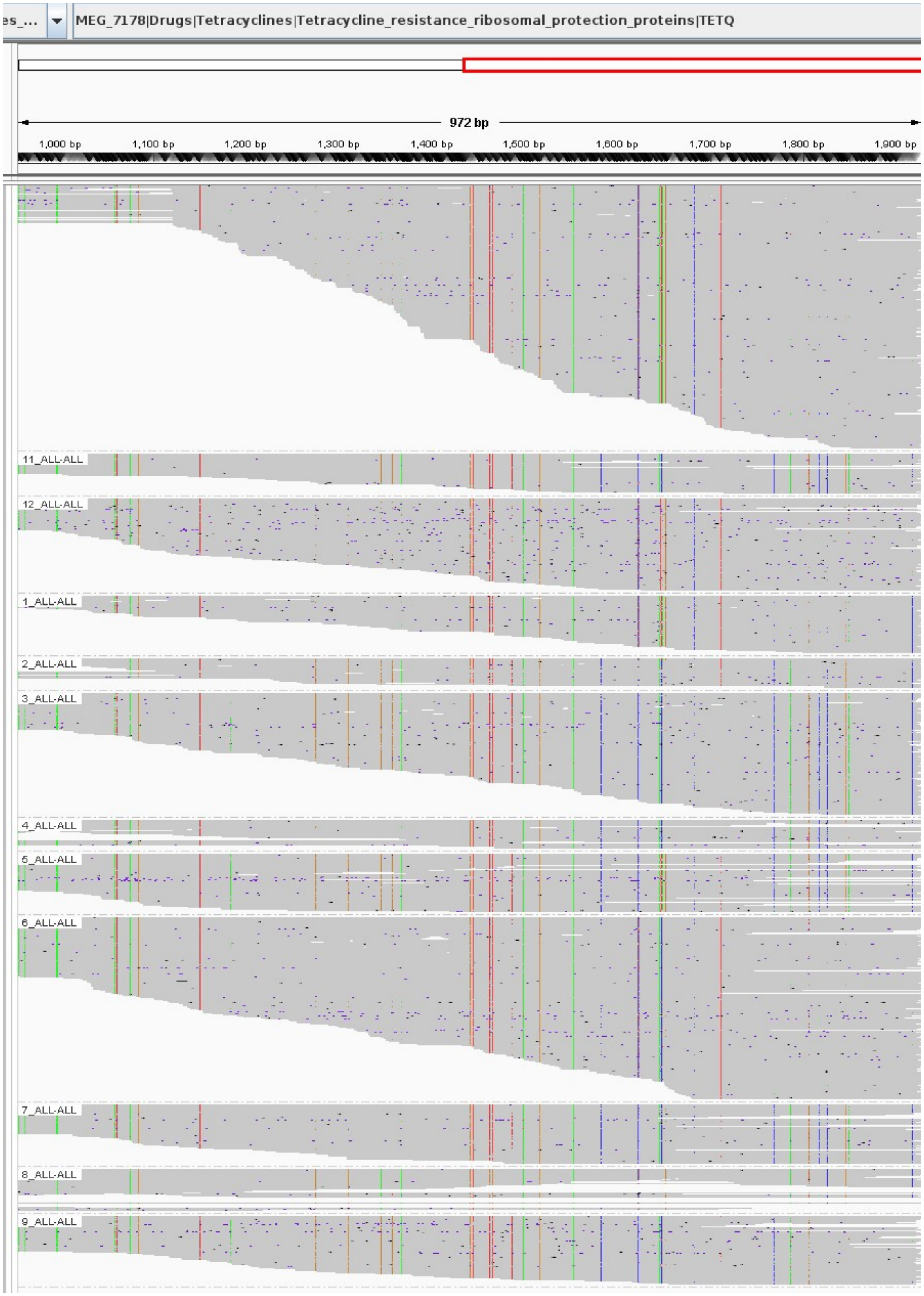


**Supplementary Figure 10.** Reads supporting a subset of the tet(Q) haplotypes shown in Fig. 4 c. Reads were downsampled to 1000x coverage per 1000 bp window for visualization purposes. Only the last 972 bp of the alignments are shown; Fig. 4 c also shows only the last 972 bp.

### References

1. Gao, Y. *et al.* abPOA: An SIMD-based C library for fast partial order alignment using adaptive band. *Bioinformatics* **37**, 2209–2211 (2021).
